# Belemnite phylogeny and decline during the mid-Cretaceous

**DOI:** 10.1101/2021.10.11.463885

**Authors:** Kevin Stevens

**Affiliations:** Ruhr-Universität Bochum, Universitätsstr. 150, 44801 Bochum

## Abstract

Belemnites are common fossil coleoid cephalopods of the Mesozoic. They began to diversify in the Triassic-Early Jurassic and maintained this diversity until the early Early Cretaceous. During the mid-Cretaceous, they declined in diversity and distribution, being restricted to only the Boreal and Austral Realm since the Turonian. Here, I present the first cladistic analysis of belemnite phylogeny, spanning taxa representative of the whole diversity and stratigraphic range of the group. This analysis shows that the usually applied subdivision of all belemnites into “Belemnitina” and “Belemnopseina” is not supported. A newly identified clade, the Pseudoalveolata, is suggested here. Pseudoalveolate belemnites represent the last remaining belemnites after the Aptian. Oceanic anoxia and warming are likely the main cause of the mid-Cretaceous belemnite decline, resulting in the Aptian-Albian dominance of the warm-adapted pseudoalveolate genus *Neohibolites*. The rise of teleost fish diversity during the mid-Cretaceous is discussed and its relevance for belemnite evolution. Some teleosts (e.g., *Enchodus*) might have taken over the mesopredator niches left by belemnites during the mid-Cretaceous, being better adapted to warming seas. Belemnites were not able to recover their earlier widespread distribution and diversity and the last remaining, disjunctly distributed families, the northern Belemnitellidae and southern Dimitobelidae, became extinct at the K/Pg-boundary.

## Introduction

Belemnites (Belemnitida) are an extinct group of stem-decabrachian coleoids (e.g., Fuchs et al., 2013; Hoffmann & Stevens, 2020). They are characterized by a calcitic rostrum, which is by far the most commonly preserved part of their internal shell. In this paper, I use the terms belemnites and Belemnitida only for these calcite-rostrum bearing species. This definition excludes groups like the Belemnoteuthida and Diplobelida, which are sometimes also referred to as belemnites. The paraphyletic assemblage of Belemnitida, Aulacoceratida, Belemnoteuthida, and Diplobelida is referred to as “belemnoids” in lieu of a proper understanding of their interrelationships at present (following Hoffmann & Stevens, 2020). Diplobelida probably represent close relatives of crown-Decabrachia (Fuchs, 2019; Fuchs et al., 2013).

The oldest belemnites are known from the Late Triassic (Carnian) of the northeastern Tethys (Zhu & Bian, 1984; Iba et al., 2012). By at least the Early Jurassic they had reached a cosmopolitan distribution and relatively high diversity, with the two usually recognized belemnite subgroups, the “Belemnitina” and “Belemnopseina” (Jeletzky, 1966), already present (e.g., Iba et al., 2014a; 2014b; Weis et al., 2015a). Belemnites continued to be diverse during the Jurassic and early Early Cretaceous (Iba et al., 2012; 2014a; 2014b).

After a major faunal and floral turnover at the Barremian-Aptian boundary (Mutterlose, 1998), the cosmopolitan belemnite genus *Neohibolites* dominated the Aptian-Albian of the Tethys, North Pacific, Austral, and Boreal Realms. *Neohibolites* went extinct during the Albian in the North Pacific Realm resulting in the local extinction of belemnites in this region (Iba et al., 2011). In the Tethys, the genus *Parahibolites* went extinct during the early Cenomanian, while *Neohibolites* went extinct during the middle Cenomanian, which led to the regional extinction of belemnites in the Tethys (Combémorel et al., 1981). The mid-Cretaceous is regarded as the time of the highest global temperatures as well as the highest global sea-level of the younger Phanerozoic, especially during the Cenomanian-Turonian (e.g., Miller et al., 2005; O’Brien et al., 2017).

*Neohibolites* is considered to be closely related to the last two surviving belemnite families: The Belemnitellidae, which were largely restricted to the Boreal Realm (Cenomanian – Maastrichtian) and the Dimitobelidae of the Austral Realm (Aptian—Maastrichtian). Only one genus of belemnitellid belemnites (*Praeactinocamax*) is present in the upper Cenomanian of the Boreal Realm, only afterwards does the group diversify (Combémorel et al, 1981; Christensen, 1997). From the Aptian until the Cenomanian, the Dimitobelidae were widespread in the southern seas. They are recorded from Aptian – Cenomanian sediments of Antarctica, India, New Guinea, Australia, southern Argentina, and New Zealand, while being restricted to Antarctica, southern Argentina, Australia, and New Zealand during the Turonian – Maastrichtian. Dimitobelids are very rare in the upper Maastrichtian and probably already became extinct before the Cretaceous/Paleogene-boundary event (Crame et al., 1996).

Several authors have discussed the ecological relationship and co-evolution of cephalopods (specifically coleoids) with teleost fishes (Packard, 1972; O’Dor & Webber, 1986; Tanner et al., 2017). The earliest teleost fishes are known from the Middle-Late Triassic boundary (Arratia, 2014). Teleost fishes diversified during the mid-Cretaceous owing in part to the origin of the hyperdiverse spiny-rayed teleosts or Acanthomorpha in the Cenomanian (e.g., Patterson, 1993; Davesne et al., 2014; Cantalice et al., 2021). Acanthomorphs are characterized by sharp spines in their fins as well as highly protrusible mouths. Teleost and especially acanthomorph diversity increased sharply after the Cretaceous/Paleogene extinction, with the morphological disparity of this group also increasing (Patterson, 1993; Friedman, 2010). Acanthomorphs represent more than half of all living teleost fishes and 25% of all extant vertebrate species (e.g., Friedman et al., 2010; Near et al., 2012; Nelson et al., 2016; Dornburg & Near, 2021).

The mid-Cretaceous was a pivotal time of belemnite evolution. This paper presents the first cladistic study of belemnite phylogeny, which identifies the Aptian as turning point in their evolution. I introduce a new monophyletic group of belemnites, the Pseudoalveolata, and point out problems with the dichotomous subdivision of all belemnites into Belemnitina and Belemnopseina. I discuss the influence of environmental factors on the belemnite decline of the mid-Cretaceous and the potential co-evolution with teleost fishes during this time.

## Methods

20 belemnite genera (*Acrocoelites, Acroteuthis, Aulacoteuthis, Belemnitella, Belemnopsis, Calabribelus, Dicoelites, Dimitobelus, Duvalia, Hibolithes, Holcobelus, Lissajousibelus, Megateuthis, Mesohibolites, Neohibolites, Oxyteuthis, Passaloteuthis, Praeactinocamax, Sinobelemnites, Schwegleria*), representative of the stratigraphic range, biogeography, and diversity of the whole group, were coded for 18 rostrum characters (Supplementary files 1, 2). Two aulacoceratid genera (*Aulacoceras, Atractites*) were also included with *Aulacoceras* serving as the outgroup for the analysis, following the hypothesis of aulacoceratid paraphyly by Keupp and Fuchs (2014). Although other fossil “belemnoid” coleoid groups are likely more closely related to the Belemnitida than aulacoceratids (e.g., Diplobelida, Belemnoteuthida), these do not have proper rostra (*sensu* Fuchs, 2012) and so do not contribute to the resolution of internal relationships of belemnites, whose phylogeny is here inferred based on rostrum characters only. Morphological data comes from several published sources (e.g., Stolley, 1911; Stoyanova-Vergilova, 1970; Combémorel, 1973; Zhu & Bian, 1984; Mutterlose, 1983; Mutterlose et al., 1987; Doyle, 1987; Christensen, 1997; Schlegelmilch, 1998; Weis et al., 2012; 2015b; Mariotti et al., 2021) and own observations.

The phylogenetic analysis was performed with TNT version 1.5 (Goloboff & Catalano, 2016). Coding practice follows suggestions of Brazeau (2011) for morphological character coding. Terminology of belemnite morphology follows Hoffmann & Stevens (2020). Exact enumeration was applied to search for most parsimonious trees (MPTs). Bootstrap support was calculated as standard (sample with replacement) with 100 replicates and jackknife support with 50% removal probability with 100 replicates, both using “traditional” tree search method. The consensus tree was calculated as a strict (Nelson) consensus.

## Results

Five MPTs were found (Fig. 1) (consistency index = 0.75, retention index = 0.8696). In all five trees Belemnitida is found as monophyletic with the aulacoceratid genus *Atractites* as sister. The resulting strict consensus tree finds a polytomy for the intrarelationships of Belemnitida. The only clades present in the strict consensus tree are clade 1 (*Aulacoteuthis, Acroteuthis*), clade 2 (*Lissajousibelus, Megateuthis, Acrocoelites, Passaloteuthis*), and clade 3 (*Dimitobelus, Neohibolites, Mesohibolites, Hibolithes*, (*Belemnitella, Praeactinocamax*)) (Fig. 2). The clade Belemnitida is well supported in the present analysis, while many taxa inside the Belemnitida remain unresolved. The here inferred subclades of the Belemnitida have relatively low support by bootstrap and jackknife values, but support for clade 3 is relatively high (63/51; Fig. 2).

**Figure 1:**
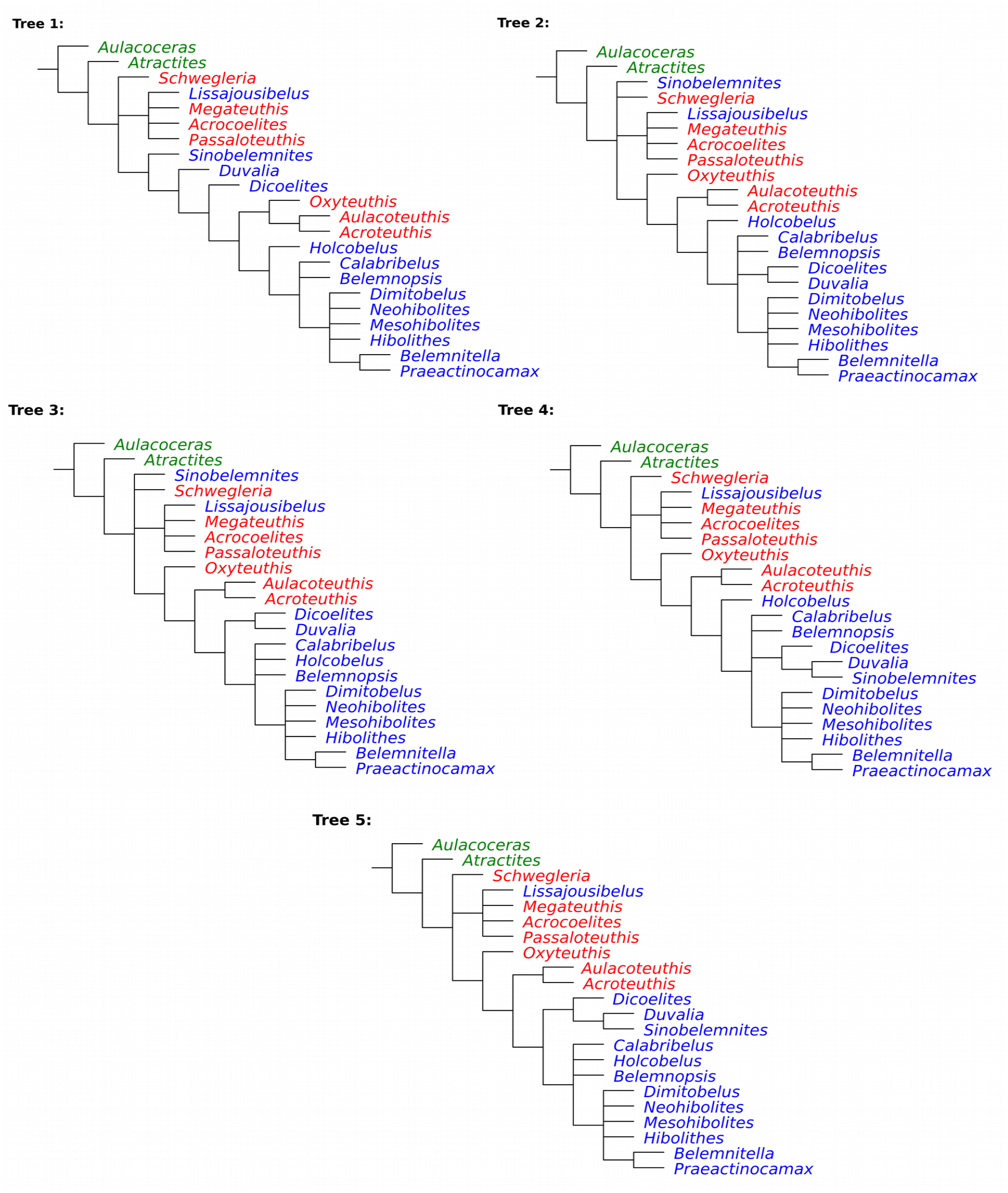
Five most parsimonious trees (MPTs) found. Aulacoceratids are in green, belemnites usually referred to the Belemnitina in red, to the Belemnopseina in blue.

**Figure 2.**
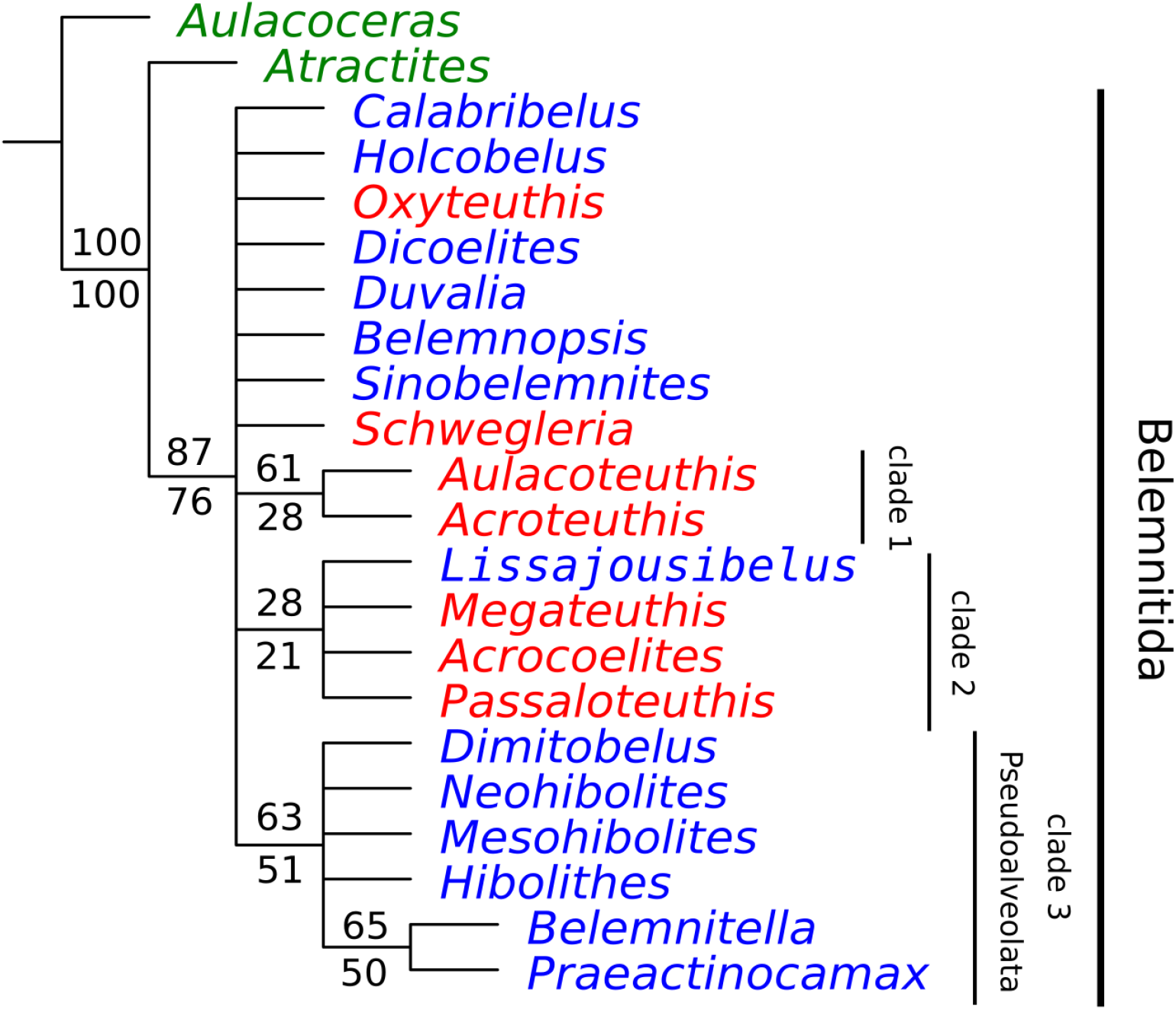
Strict consensus tree of five MPTs. Aulacoceratids are in green, belemnites usually referred to the Belemnitina in red, to the Belemnopseina in blue. Bootstrap and jackknife support values are indicated above and below nodes, respectively.

## Discussion

### Belemnite phylogeny

Since Jeletzky (1966), belemnites were usually subdivided into two suborders; Belemnitina and Belemnopseina, with some members of the former seen as ancestral to the latter. Belemnitina groups taxa with apical furrows and Belemnopseina taxa with alveolar furrows. It has long been known that many belemnites do not fit into this dichotomous subdivision. For example, the genus *Lissajousibelus* displays a ventral furrow with slit in addition to dorsolateral apical furrows but was considered close to “Belemnopseina” by Weis et al. (2015b). In the present analysis it was found to be closely related to typical “Belemnitina”, *Passaloteuthis, Acrocoelites*, and *Megateuthis*. The Duvaliidae, Dicoelitidae, and Holcobelidae have also been of uncertain phylogenetic placement (e.g., Stevens, 1964; Weis et al., 2012). The present analysis unfortunately cannot resolve these taxa with certainty either, but in four of the five MPTs *Duvalia* and *Dicoelites* form a clade (Fig. 4), which might give impetus for further analyses of a close relationship. The two holcobelids *Holcobelus* and *Calabribelus*, while not found to form a monophylum in any of the MPTs, are close to “belemnopseine” belemnites (Fig. 1), confirming Weis et al. (2012).

**Figure 3:**
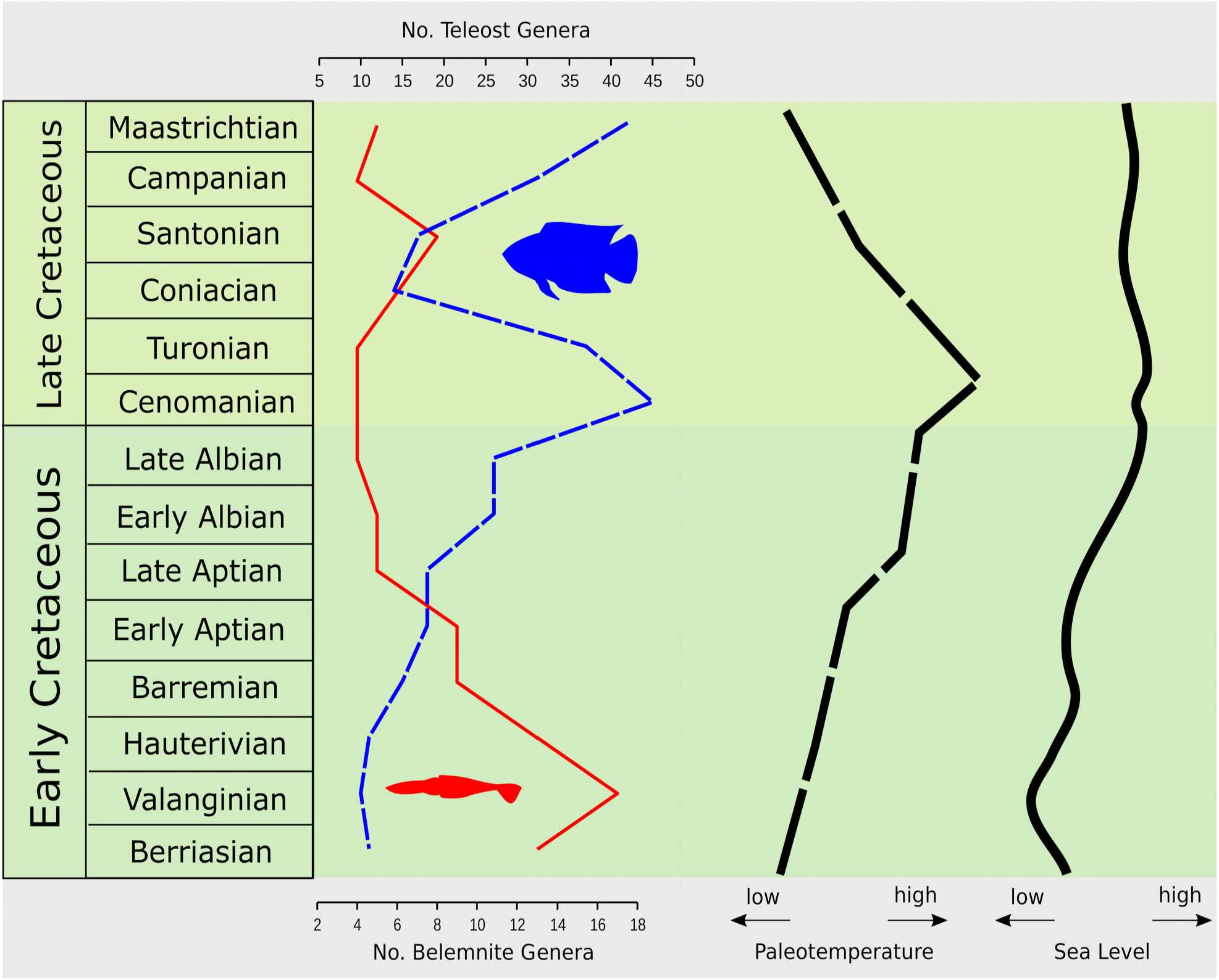
Genus richness of belemnites and teleosts with paleotemperature and sea level. Diversity data for belemnites from Mutterlose (1988), Doyle (1992), Christensen (1997), Williamson & Henderson (2015), and for teleosts from the Paleobiology Database (hps://paleobiodb.org/navigator/#/7b512dac). Simplified paleotemperature and eustatic seawater-level curves based on O’Brien et al. (2017) and Haq (2014).

**Figure 4:**
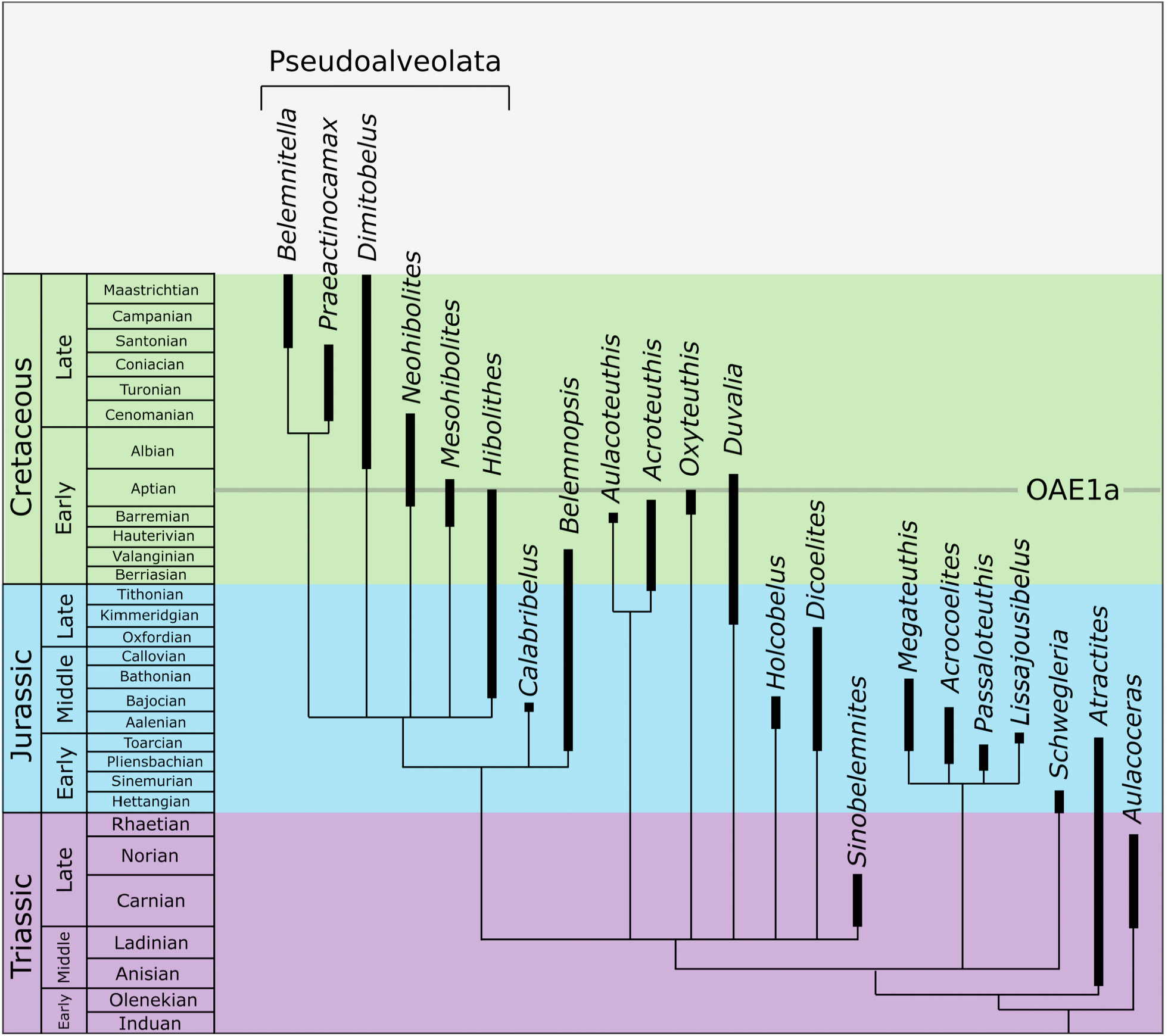
Combinable components consensus tree with stratigraphic ranges of the analyzed aulacoceratid and belemnite genera (Stevens, 1964; Combémorel et al., 1981; Mutterlose, 1983; 1988; 1998; Doyle, 1987; Christensen, 1997; Schlegelmilch, 1998; Iba et al., 2012; Weis et al., 2012, 2015; Rogov et al., 2019; Mariotti et al., 2021).

The oxyteuthid *Aulacoteuthis* and cylindroteuthid *Acroteuthis* are sister taxa in the present analysis (Fig. 2). While no monophyletic Oxyteuthidae have been found, the present analysis might point towards a close relationship of the oxyteuthids to *Acroteuthis* as suggested by Mutterlose (1983). Still, the problem of the phylogenetic placement and evolution of the Oxyteuthidae needs further focused analyses, especially with respect to the oxyteuthids of the Russian platform (Mutterlose & Baraboshkin, 2003; Baraboshkin & Mutterlose, 2004).

Overall, there is no support from the present analysis for a dichotomous subdivision of all belemnites into Belemnitina and Belemnopseina, as these groups are usually defined (Figs. 1, 2). Based on the present analysis, all belemnites cannot be grouped under the traditional simple dichotomous scheme, although a phylogenetic redefinition of Belemnitina and Belemnopseina seems possible. The present consensus topology represents only a first step towards a better resolved phylogeny of all belemnites. To achieve further resolution, it will likely be necessary to detect and evaluate further microstructural and geochemical data of several belemnite taxa, in order to get a better understanding of the phylogeny of this group.

A name is suggested here for the newly identified clade 3 as the unranked Pseudoalveolata (Fig. 2). Pseudoalveolata is defined by the synapomorphy of pseudoalveolus formation, i.e., the alveolus of members of this group is usually found to be in part eroded. A pseudoalveolus might be present as a concave secondary deepening of the alveolus (e.g., some *Dimitobelus*) or as formation of a conical structure by complete loss of the rostrum cavum and anterior rostrum solidum (e.g., *Hibolithes, Praeactinocamax*). Pseudoalveolus types have been of great importance in the phylogeny of the Belemnitellidae (e.g., Košťak, 2012). Contrary to suggestions by Dauphin et al. (2007) and Košťak & Wiese (2008), there is no conclusive evidence for the alveolus of pseudoalveolate belemnites being of primarily aragonitic composition, it instead consisted of calcite with primarily high organic contents (Stevens, 2017; Stevens et al., 2017). Inside the Pseudoalveolata the belemnitellids *Praeactinocamax* and *Belemnitella* were found as sister taxa, but a more extensive analysis of the Late Cretaceous family Belemnitellidae would be needed to confirm its monophyly (Fig. 2).

### Belemnite decline during the mid-Cretaceous

Belemnite diversity was significantly reduced during the Aptian (Fig. 3). In the Albian, only members of the Pseudoalveolata persist (Fig. 4), namely the genera *Neohibolites* and *Parahibolites* in the Tethys and Boreal Realm as well as the Dimitobelidae in the Austral Realm (e.g., Christensen, 1997; Doyle, 1992). The early Aptian is characterized by OAE1a, an oceanic anoxic event triggered by volcanism of the Ontong-Java Plateau (e.g., Jenkyns, 2010; Erba et al., 2015). Volcanism resulted in a pronounced warming and rise of oxygen minimum zones in the oceans, resulting in widespread deposition of organic rich sediments and the disappearance of metazoan-dominated carbonate platforms in the Tethys (Bottini & Mutterlose, 2012; Huck et al., 2014; Hueter et al., 2019; Bauer et al., 2021).

OAE1a was likely the driving factor in the extinction of the oxyteuthids, which are usually viewed as the last members of the “Belemnitina”. Members of the last-occurring genus *Oxyteuthis* are recorded from German and Russian sections until black shale formation in association with OAE1a (Hoffmann & Mutterlose, 2011; Lehmann et al., 2012; Rogov et al., 2019). The pseudoalveolate *Neohibolites* originated in the earliest Aptian and is regarded as a warm-adapted genus (Mutterlose, 1998). Its survival might be linked to being better able to cope warming and anoxia in association with OAE1a. The local extinction of *Neohibolites* during the late Albian in the North Pacific Realm is thought to be linked to a cooling phase in this region (Iba et al., 2011) and similarly, warm-water *Neohibolites* might have become extinct due to cooling during the middle Cenomanian Plenus Cold Event (Combémorel et al., 1981; Ernst et al., 1983). The Belemnitellidae originated in the early Cenomanian with *Praeactinocamax* and were apparently well-adapted to cooler conditions, which is suggested by the mass-occurrence of *Praeactinocamax plenus* during the short-lived mid-Cenomanian Plenus Cold Event, when they even spread into the northern Tethys (Gale & Christensen, 1996). During the Aptian there are two phases during which belemnites of the mostly Tethyan family Duvaliidae spread northward, shortly after OAE1a and during the late Aptian (Mutterlose, 1998). Duvaliids become extinct later in the Aptian (Fig. 4) and therefore represent the last surviving non-pseudoalveolate belemnites. Dimitobelids also seem to have been adapted to cool conditions. From the Turonian onwards they are restricted to the shelf seas of the southernmost remnants of Gondwana (Doyle, 1992; Crame et al., 1996), potentially becoming extinct further north due to warming during the mid-Cretaceous. While pseudoalveolate belemnites form a monophyletic group, they do not seem to have shared the same ecological preferences regarding temperature.

Although belemnites are often viewed as nektobenthic, this assumption is problematic (Hoffmann & Stevens, 2020). A demersal habitat of several belemnites is unlikely given their common occurrence in sediments deposited below anoxic bottom waters but cannot be excluded for all belemnite taxa (Rexfort & Mutterlose, 2006; Mutterlose et al., 2010; Stevens et al., 2014). It therefore seems unlikely that anoxia affected belemnites directly in their upper water habitat but warming and the general effects on ecosystems of the spread of anoxic waters might have resulted in their decline during the mid-Cretaceous.

Molecular clocks indicate a diversification of teleosts during the mid-Cretaceous, which is congruent with fossil data based on the skeletal (Fig. 3, Cavin et al., 2007; Near et al., 2012; Guinot & Cavin, 2016) and otolith record (Schwarzhans, 2018). This radiation has been shown not to be an artifact of a “lagerstätten effect” (Cavin & Forey, 2007). It seems unlikely that the newly emerged acanthomorphs and other teleosts replaced belemnites via competitive displacement. Competitive displacement in the fossil record can be shown, if: 1) both groups show an inverse relationship of their respective diversity, 2) they can be regarded to have utilized the same resources, and 3) both groups evolved in allopatry before dispersal of one group into the other’s range (Krause, 1986). All those points cannot be conclusively shown for teleosts and belemnites. Although both groups show largely inverse diversity trends, low recorded diversity of teleosts from the Coniacian to Campanian (Fig. 3)is likely a sampling artifact (Guinot & Cavin, 2016). Belemnites were mesopredators, likely being restricted to a carnivorous diet (Hoffmann & Stevens, 2020), while teleosts during the Cretaceous display diverse feeding strategies (e.g., herbivores; Davesne et al., 2018). Both groups certainly did not evolve allopatrically, teleosts and belemnites are both known from the Triassic onward in the marine realm. It seems more plausible that higher eustatic sea level as well as high temperatures favored the radiation of teleosts during the mid-Cretaceous (Cavin et al., 2007) and they diversified into some of the niches earlier occupied by belemnite species. For example, the widespread aulopiform family Enchodontidae were medium-sized mesopredatory fishes. The often highly abundant type genus *Enchodus* originated in the latest Albian/Cenomanian and was a very common component of ichthyofaunas until the end of the Cretaceous (e.g., Fielitz & González-Rodríguez, 2010). A faunal replacement of belemnites by crown-octobrachian and crown-decabracian coleoids likely also occurred during the mid-Cretaceous (Iba et al., 2011; Tanner et al., 2017).

The Cenomanian decline of belemnites has been suggested to be the main factor in the extinction of ichthyosaurs at the Cenomanian/Turonian boundary (Bardet, 1992). A similar belemnite decline during the Toarcian was also suggested to have affected ichthyosaurs during this time (Martin et al., 2012). Later studies, however, argued for an environmentally driven extinction of ichthyosaurs during OAE2 (e.g., Fischer et al., 2016). A new appreciation of Cretaceous ichthyosaur diversity and disparity also argues against an extinction of ichthyosaurs due to the decline of belemnites. Several of the last ichthyosaurs belonged to the apex predator or generalist niche and were not restricted to soft-bodied prey like belemnites (e.g., Fischer, 2016).

## Conclusions

The first cladistic phylogeny of the belemnites (Belemnitida) is presented. Belemnitida as a monophyletic group is well supported but the usually applied dichotomous subdivision of all belemnites into Belemnitina and Belemnopseina finds no support. A new subgroup, consisting of a subset of the “belemnopseine” belemnites, the unranked Pseudoalveolata, is suggested here.

Belemnite diversity was reduced sharply during the Aptian, afterwards only belemnites of the Pseudoalvolata remained. In the late Albian and Cenomanian, the geographic distribution of belemnites was reduced, after the middle Cenomanian the group persists only as the Belemnitellidae of the Boreal Realm and Dimitobelidae of the Austral Realm. High sea-surface temperatures, high sea-level, and phases of widespread anoxia during the mid-Cretaceous seem to have resulted in their extinction and reduced distribution and diversity. The radiation of teleost fishes during the mid-Cretaceous cannot be causally linked to the decline of belemnites but some of the vacant mesopredator niches of belemnites might have been filled by teleosts.

Belemnites, because of their high fossilization potential and often high abundance in the marine fossil record, might be of key importance in tracking faunal changes of pelagic realm of the Jurassic and Cretaceous. The present study is only a first step and a phylogeny based on more taxa and characters is needed to further resolve their evolutionary history and paleoecology.

## Acknowledgments

The Willi Hennig Society is acknowledged for providing TNT. I am thankful to René Hoffmann for discussions and remarks on an earlier version of this manuscript.

## References

Arratia, G., 2013. Morphology, taxonomy, and phylogeny of Triassic pholidophorid fishes (Actinopterygii, Teleostei). Journal of Vertebrate Paleontology 33, 1–138. doi:10.1080/02724634.2013.835642

Bauer, K.W., Bottini, C., Frei, R., Asael, D., Planavsky, N.J., Francois, R., McKenzie, N.R., Erba, E., Crowe, S.A., 2021. Pulsed volcanism and rapid oceanic deoxygenation during Oceanic Anoxic Event 1a. Geology. https://doi.org/10.1130/G49065.1

Baraboshkin, E.J., Mutterlose, J., 2004. Correlation of the Barremian belemnite successions of northwest Europe and the Ulyanovsk – Saratov area (Russian Platform). Acta Geologica Polonica 54, 499–510.

Bardet, N., 1992. Stratigraphic evidence for the extinction of the ichthyosaurs. Terra Nova 4, 649–656.

Bottini, C., Mutterlose, J., 2012. Integrated stratigraphy of Early Aptian black shales in the Boreal Realm: calcareous nannofossil and stable isotope evidence for global and regional processes. nos 45, 115–137. https://doi.org/10.1127/0078-0421/2012/0017

Brazeau, M.D., 2011. Problematic character coding methods in morphology and their effects. Biological Journal of the Linnean Society 104, 489–498. https://doi.org/10.1111/j.1095-8312.2011.01755.x

Cantalice, K.M., Than-Marchese, B.A., Villalobos-Segura, E., 2021. A new Cenomanian acanthomorph fish from the El Chango quarry (Chiapas, south-eastern Mexico) and its implications for the early diversification and evolutionary trends of acanthopterygians. Papers in Palaeontology 7, 1699–1726. https://doi.org/10.1002/spp2.1359

Cavin, L., Forey, P.L., 2007. Using ghost lineages to identify diversification events in the fossil record. Biology Letters 3, 201–204. doi:10.1098/rsbl.2006.0602

Cavin, L., Forey, P.L., Lécuyer, C., 2007. Correlation between environment and Late Mesozoic ray-finned fish evolution. Palaeogeography, Palaeoclimatology, Palaeoecology 245, 353– 367. doi:10.1016/j.palaeo.2006.08.010

Christensen, W.K., 1997. The Late Cretaceous belemnite family Belemnitellidae: Taxonomy and evolutionary history. Bulletin of the Geological Society of Denmark 44, 59–88.

Christensen, W.K., 2002. Palaeobiology, phylogeny and palaeobiogeography of belemnoids and related coleoids. Berliner Paläobiologische Abhandlungen 1, 18–21.

Combémorel, R., 1973. Les Duvaliidae (Pavlow) du cretace inferieur francais. Document des laboratoires de geology de la faculte des sciences de Lyon 57, 131–186.

Combémorel, R., Christensen, W.K., Naidin, D.P., Spaeth, C., 1981. Les Bélemnites. Cretaceous Research 2, 283–286.

Crame, J.A., Lomas, S.A., Pirrie, D., Luther, A., 1996. Late Cretaceous extinction patterns in Antarctica. Journal of the Geological Society, London 153, 503–506.

Dauphin, Y., Williams, C.T., Barskov, I.S., 2007. Aragonitic rostra of the Turonian belemnitid Goniocamax: Arguments from diagenesis. Acta Palaeontologica Polonica 52, 85.

Davesne, D., Gueriau, P., Dutheil, D.B., Bertrand, L., 2018. Exceptional preservation of a Cretaceous intestine provides a glimpse of the early ecological diversity of spiny-rayed fishes (Acanthomorpha, Teleostei). Scientific Reports 8. https://doi.org/10.1038/s41598-018-26744-3

Dornburg, A., Near, T.J., 2021. The Emerging Phylogenetic Perspective on the Evolution of Actinopterygian Fishes. Annual Reviews in Ecology and Evolution. Syst. 52, annurev-ecolsys-122120-122554. https://doi.org/10.1146/annurev-ecolsys-122120-122554

Doyle, P., 1987. The Cretaceous Dimitobelidae (Belemnitida) of the Antarctic Peninsula region. Palaeontology 30, 147–177.

Doyle, P., 1992. A review of the biogeography of cretaceous belemnites. Palaeogeography, Palaeoclimatology, Palaeoecology 92, 207–216. https://doi.org/10.1016/0031-0182(92)90082-G

Erba, E., Duncan, R.A., Bottini, C., Tiraboschi, D., Weissert, H., Jenkyns, H.C., Malinverno, A., 2015. Environmental consequences of Ontong Java Plateau and Kerguelen Plateau volcanism. Geological Society of America Special Papers 511, SPE511–15.

Ernst, G., Schmid, F., Seibertz, E., 1983. Event-Stratigraphie im Cenoman und Turon von NW-Deutschland. Zitteliana 19, 531–554.

Fielitz, C., González-Rodríguez, K.A., 2010. A new species of Enchodus (Aulopiformes: Enchodontidae) from the Cretaceous (Albian to Cenomanian) of Zimapán, Hidalgo, México. Journal of Vertebrate Paleontology 30, 1343–1351. https://doi.org/10.1080/02724634.2010.501438

Fischer, V., 2016. Taxonomy of Platypterygius campylodon and the diversity of the last ichthyosaurs. PeerJ 4, e2604. https://doi.org/10.7717/peerj.2604

Fischer, V., Bardet, N., Benson, R.B.J., Arkhangelsky, M.S., Friedman, M., 2016. Extinction of fish-shaped marine reptiles associated with reduced evolutionary rates and global environmental volatility. Nature Communications 7, 10825. https://doi.org/10.1038/ncomms10825

Friedman, M., 2010. Explosive morphological diversification of spiny-finned teleost fishes in the aftermath of the end-Cretaceous extinction. Proceedings of the Royal Society B: Biological Sciences 277, 1675–1683. doi:10.1098/rspb.2009.2177

Friedman, M., Keck, B.P., Dornburg, A., Eytan, R.I., Martin, C.H., Hulsey, C.D., Wainwright, P.C., Near, T.J., 2013. Molecular and fossil evidence place the origin of cichlid fishes long after Gondwanan rifting. Proceedings of the Royal Society B: Biological Sciences 280, 20131733–20131733. doi:10.1098/rspb.2013.1733

Fuchs, D., 2012. The “rostrum”-problem in coleoid terminology – an attempt to clarify inconsistencies. Geobios 45, 29–39. https://doi.org/10.1016/j.geobios.2011.11.014

Fuchs, D., 2019. Part M, Coleoidea, Chapter 23E: Systematic descriptions: Diplobelida. Treatise Online 118.

Gale, A.S., Christensen, W.K., 1996. Bulletin of the Geological Society of Denmark 43, 68–77.

Goloboff, P.A., Catalano, S.A., 2016. TNT version 1.5, including a full implementation of phylogenetic morphometrics. Cladistics 32, 221–238. https://doi.org/10.1111/cla.12160

Guinot, G., Cavin, L., 2016. ‘Fish’ (Actinopterygii and Elasmobranchii) diversification patterns through deep time: ‘Fish’ diversification patterns through deep time. Biological Reviews 91, 950–981. https://doi.org/10.1111/brv.12203

Haq, B.U., 2014. Cretaceous eustasy revisited. Global and Planetary Change 113, 44–58. https://doi.org/10.1016/j.gloplacha.2013.12.007

Hoffmann, R., Mutterlose, J., 2011. Stratigraphie und Cephalopodenfauna des Unter-Apt von Alstätte (NRW). Geologie und Paläontologie in Westfalen 80, 43–59.

Hoffmann, R., Stevens, K., 2020. The palaeobiology of belemnites – foundation for the interpretation of rostrum geochemistry. Biological Reviews 95, 94–123. https://doi.org/10.1111/brv.12557

Huck, S., Stein, M., Immenhauser, A., Skelton, P.W., Christ, N., Föllmi, K.B., Heimhofer, U., 2014. Response of proto-North Atlantic carbonate-platform ecosystems to OAE1a-related stressors. Sedimentary Geology 313, 15–31. https://doi.org/10.1016/j.sedgeo.2014.08.003

Hueter, A., Huck, S., Bodin, S., Heimhofer, U., Weyer, S., Jochum, K.P., Immenhauser, A., 2019. Central Tethyan platform-top hypoxia during Oceanic Anoxic Event 1a. Climate of the Past 15, 1327–1344. https://doi.org/10.5194/cp-15-1327-2019

Iba, Y., Mutterlose, J., Tanabe, K., Sano, S., Misaki, A., Terabe, K., 2011. Belemnite extinction and the origin of modern cephalopods 35 m.y. prior to the Cretaceous-Paleogene event. Geology 39, 483–486.

Iba, Y., Sano, S., Mutterlose, J., Kondo, Y., 2012. Belemnites originated in the Triassic — A new look at an old group. Geology 40, 911–914. doi:10.1130/G33402.1

Iba, Y., Sano, S., Mutterlose, J., 2014a. The Early Evolutionary History of Belemnites: New Data from Japan. PLOS ONE 9, e95632.

Iba, Y., Sano, S., Rao, X., Fuchs, D., Chen, T., Weis, R., Sha, J., 2014b. Early Jurassic belemnites from the Gondwana margin of the Southern Hemisphere–Sinemurian record from South Tibet. Gondwana Research. doi:10.1016/j.gr.2014.06.007

Jeletzky, J.A., 1966. Comparative morphology, phylogeny, and classification of fossil Coleoidea. The University of Kansas Paleontological Contributions 7, 162 pp.

Jenkyns, H.C., 2010. Geochemistry of oceanic anoxic events. Geochemistry, Geophysics, Geosystems. 11, n/a-n/a. https://doi.org/10.1029/2009GC002788

Keupp, H., Fuchs, D., 2014. Different regeneration mechanisms in the rostra of aulacocerids (Coleoidea) and their phylogenetic implications. Göttingen Contributions to Geosciences 77, 13–20.

Košťák, M., 2012. On the Turonian origin of the Goniocamax-Belemnitella stock (Cephalopoda, Coleoidea). Geobios 45, 79–85. https://doi.org/10.1016/j.geobios.2011.11.004

Košťák, M., Wiese, F., 2008. Lower Turonian record of belemnite Praeactinocamax from NW Siberia and its palaeogeographic significance. Acta Palaeontologica Polonica 53, 669–678.

Krause, D.W., 1986. Competitive exclusion and taxonomic displacement in the fossil record; the case of rodents and multituberculates in North America. Rocky Mountain Geology 24, 95–117.

Lehmann, J., Friedrich, O., Von Bargen, D., Hemker, T., 2012. Early Aptian bay deposits at the southern margin of the Lower Saxony Basin: Integrated Stratigraphy, palaeoenvironment and OAE1a. Acta Geologica Polonica 62, 35–62. https://doi.org/10.2478/v10263-012-0002-2

Mariotti, N., Pignatti, J., Riegraf, W. 2021. Part M, Coleoidea, Chapter 23E: Systematic descriptions: Diplobelida. Treatise Online 148.

Martin, J.E., Fischer, V., Vincent, P., Suan, G., 2012. A longirostrine Temnodontosaurus (Ichthyosauria) with comments on Early Jurassic ichthyosaur niche partitioning and disparity. Palaeontology 55, 995–1005. doi:10.1111/j.1475-4983.2012.01159.x

Matschiner, M., 2019. Gondwanan vicariance or trans-Atlantic dispersal of cichlid fishes: a review of the molecular evidence. Hydrobiologia 832, 9–37. https://doi.org/10.1007/s10750-018-3686-9

Miller, K.G., Kominz, M.A., Browning, J.V., Wright, J.D., Mountain, G.S., Katz, M.E., Sugarman, P.J., Cramer, P.S., Christie-Blick, N., Pekar, S.F., 2005. The Phanerozoic record of global sea-level change. Science 310, 1293–1298.

Mutterlose, J. 1983. Phylogenie und Biostratigraphie der Unterfamilie Oxyteuthinae (Belemnitida) aus dem Barreme (Unter-Kreide) NW-Europas. Palaeontographica Abt. A 180, 1–90.

Mutterlose, J., 1988. Migration and Evolution Patterns in Upper Jurassic and Lower Cretaceous Belemnites. In: Wiedmann, J. & Kullmann, J. (eds.), Cephalopods – Present and Past, Schweizerbart’sche Verlagsbuchhandlung, 525–537.

Mutterlose, J., 1998. The Barremian–Aptian turnover of biota in northwestern Europe: evidence from belemnites. Palaeogeography, Palaeoclimatology, Palaeoecology 144, 161– 173.

Mutterlose, J., Baraboshkin, E.J., 2003. Taxonomy of the Early Cretaceous belemnite species Aulacoteuthis absolutiformis (Sinzow, 1877) and its type status. Berliner Paläobiologische Abhandlungen 3, 179–187.

Mutterlose, J., Pinckney, G., Rawson, P.F., 1987. The belemnite Acroteuthis in the Hibolites beds (Hauterivian-Barremian) of north-west Europe. Palaeontology 30, 635–645.

Near, T.J., Eytan, R.I., Dornburg, A., Kuhn, K.L., Moore, J.A., Davis, M.P., Wainwright, P.C., Friedman, M., Smith, W.L., 2012. Resolution of ray-finned fish phylogeny and timing of diversification. Proceedings of the National Academy of Sciences 109, 13698–13703. https://doi.org/10.1073/pnas.1206625109

Nelson, J. S., Grande, T. C., Wilson, M. V. 2016. Fishes of the World. John Wiley & Sons.

O’Brien, C.L., Robinson, S.A., Pancost, R.D., Sinninghe Damsté, J.S., Schouten, S., Lunt, D.J., Alsenz, H., Bornemann, A., Bottini, C., Brassell, S.C., Farnsworth, A., Forster, A., Huber, B.T., Inglis, G.N., Jenkyns, H.C., Linnert, C., Littler, K., Markwick, P., McAnena, A., Mutterlose, J., Naafs, B.D.A., Püttmann, W., Sluijs, A., van Helmond, N.A.G.M., Vellekoop, J., Wagner, T., Wrobel, N.E., 2017. Cretaceous sea-surface temperature evolution: Constraints from TEX_86_ and planktonic foraminiferal oxygen isotopes. Earth-Science Reviews 172, 224–247. https://doi.org/10.1016/j.earscirev.2017.07.012

O’Dor, R.K., Webber, D.M., 1986. The constraints on cephalopods: why squid aren’t fish. Canadian Journal of Zoology 64, 1591–1605.

Packard, A., 1972. Cephalopods and fish: the limits of convergence. Biological Reviews 47, 241–307.

Rogov, M.A., Shchepetova, E.V., Ippolitov, A.P., Seltser, V.B., Mironenko, A.A., Pokrovsky, B.G., Desai, B.G., 2019. Response of cephalopod communities on abrupt environmental changes during the early Aptian OAE1a in the Middle Russian Sea. Cretaceous Research 96, 227– 240. https://doi.org/10.1016/j.cretres.2019.01.007

Schlegelmilch, R., 1998. Die Belemniten des suddeutschen Jura. Ein Bestimmungsbuch fur Geowissenschaftler und Fossiliensammler. Gustav Fischer Verlag, 151 pp.

Schwarzhans, W., 2018. A review of Jurassic and Early Cretaceous otoliths and the development of early morphological diversity in otoliths. Neues Jahrbuch für Geologie und Paläontologie 287, 75–121. https://doi.org/10.1127/njgpa/2018/0707

Stevens, G.R., 1964. The belemnite genera Dicoelites Boehm and Prodicoelites Stolley. Palaeontology 7, 606–620.

Stevens, K., 2017. Calcitic skeletons of recent and fossil Coleoidea: Biological, environmental, or diagenetic control? PhD thesis, Bochum. 185 pp. http://hss-opus.ub.ruhr-unibochum.de/opus4/frontdoor/index/index/docId/5374

Stevens, K., Mutterlose, J., Schweigert, G., 2014. Belemnite ecology and the environment of the Nusplingen Plattenkalk (Late Jurassic, southern Germany): evidence from stable isotope data. Lethaia n/a-n/a. https://doi.org/10.1111/let.12076

Stevens, K., Griesshaber, E., Schmahl, W., Casella, L.A., Iba, Y., Mutterlose, J., 2017. Belemnite biomineralization, development, and geochemistry: The complex rostrum of Neohibolites minimus. Palaeogeography, Palaeoclimatology, Palaeoecology 468, 388–402. https://doi.org/10.1016/j.palaeo.2016.12.022

Stolley E. 1911. Beitrage zur Kenntnis der Cephalopoden der norddeutschen unteren Kreide, I. Die Belemnitiden der norddeutschen unteren Kreide, 1. Die Belemniten des norddeutschen Gaults (Aptiens und Albiens). Geologische und Palaontologische Abhandlungen, Band 10, Heft 3.

Stoyanova-Vergilova, M., 1970. Les Fossiles de Bulgarie, IVa Cretace inferieur, Belemnitida [in Bulgarian]. Academie Bulgare des Sciences, 72 pp.

Tanner, A.R., Fuchs, D., Winkelmann, I.E., Gilbert, M.T.P., Pankey, M.S., Ribeiro, Â.M., Kocot, K.M., Halanych, K.M., Oakley, T.H., da Fonseca, R.R., Pisani, D., Vinther, J., 2017. Molecular clocks indicate turnover and diversification of modern coleoid cephalopods during the Mesozoic Marine Revolution. Proceedings of the Royal Society B: Biological Sciences 284, 20162818. https://doi.org/10.1098/rspb.2016.2818

Weis, R., Mariotti, N., Di Cencio, A., 2015a. Systematics and evolutionary implications of Early Jurassic belemnites from the Peri-Mediterranean Tethys. Paläontologische Zeitschrift. https://doi.org/10.1007/s12542-015-0265-5

Weis, R., Dzyuba, O.S., Mariotti, N., Chesnier, M., 2015b. Lissajousibelus nov. gen., an Early Jurassic canaliculate belemnite from Normandy, France. Swiss Journal of Palaeontology 134, 289–300. https://doi.org/10.1007/s13358-015-0086-x

Williamson, T., Henderson, R.A., 2015. Pumiliobelus, a new dwarf coleoid genus (Belemnoidea: Dimitobelidae) from the Cenomanian of Western Australia. Journal of Paleontology 89, 183–188. https://doi.org/10.1017/jpa.2014.15

Zhu, K.-Y., Bian, Z.-X., 1984. Sinobelemnitidae, a new family of Belemnitida from the Upper Triassic of Longmenshan, Sichuan. Acta Palaeontologica Sinica 23, 300–325.

